# Association between soil properties and Grapevine Leaf Stripe Disease (Esca Complex) symptom expression. Part I: soil physical, hydrological and chemical characteristics

**DOI:** 10.64898/2026.10.28.762510

**Authors:** Francesco Bigazzi, Simone Priori, Sara Del Duca, Francesco Vitali, Stefano Mocali, Giuseppe Carella, Laura Mugnai

**Affiliations:** University of Florence, Department of Agricultural, Food, Environmental and Forestry Science and Technology (DAGRI), Plant pathology and Entomology section, P.le delle Cascine 28, 50142 Firenze, Italy; University of Tuscia, Department of Agriculture and Forest Sciences (DAFNE), via San Camillo de Lellis, 01100 Viterbo; Council for Agricultural Research and Economics - Research Centre for Agriculture and Environment (CREA-AA), Via di Lanciola 12/A, 50125 Firenze, Italy; Council for Agricultural Research and Economics - Research Centre for Genomics and Bioinformatics (CREA-GB), Via San Protaso 302, 29017 Fiorenzuola D’Arda, Italy

**Keywords:** viticulture, pathogens, abiotic stress, soil mapping, proximal sensing

## Abstract

This study is the first of two companion papers (Part II addresses rhizosphere microbial communities) and aims to investigate the influence of soil properties on GLSD foliar symptom expression by comparing high recurrence of symptomatic vines areas (HRSVAs) and low recurrence of symptomatic vines areas (LRSVAs). The research was conducted in an approximately 1.75-hectare Cabernet Sauvignon vineyard in Tuscany, Italy. Electromagnetic induction (EMI) proximal sensing was combined with kriging-based maps of grapevine leaf stripe disease (GLSD) symptom recurrence over a five-year period, allowing the identification of four vineyard sectors representing contrasting levels of symptom recurrence (high vs. low) and soil apparent electrical conductivity (ECa; medium vs. high). One soil profile was excavated in each sector and characterised through pedological description and physicochemical analysis. GLSD symptoms showed a heterogeneous yet spatially structured distribution, with distinct zones exhibiting contrasting recurrence. Soil properties differed markedly between HRSVAs and LRSVAs. LRSVAs were characterised by clay loam to silty clay loam textures, high lime content, alkaline pH, higher available water capacity (AWC), and lower available manganese concentrations (4.5–4.8 mg kg□¹). In contrast, HRSVAs had sandy loam to sandy clay loam textures, subalkaline pH, greater soil fertility (higher nitrogen and organic matter contents), deeper root distribution, and higher available manganese concentrations (14.7–14.8 mg kg□¹). These findings highlight the importance of investigating soil physicochemical properties in relation to GLSD symptom expression, as both macro- and micronutrient availability may be associated with disease dynamics. The higher manganese availability observed in HRSVAs, together with recent evidence that manganese can stimulate the wood-degrading activity of *Fomitiporia mediterranea*, warrants further investigation into the potential association between soil manganese bioavailability and disease expression. The spatial correspondence between soil heterogeneity and GLSD symptom recurrence highlights the value of high-resolution investigations of soil–plant interactions for improving our understanding of this complex disease.

## Introduction

Grapevine trunk diseases (GTDs) represent a growing threat to viticulture worldwide (Gramaje *et al*., 2018; Moret *et al*., 2019). Among these, the syndromes grouped under the esca complex of diseases (ECDs, *sensu* Surico, 2009) are some of the most widespread and damaging in Europe, with an increasing economic impact (Bertsch *et al*., 2012; Bruez *et al*., 2021a).

The main fungal agents involved globally in ECDs are two tracheomycotic ascomycetes, *Phaeoacremonium minimum* and *Phaeomoniella chlamydospora*, and wood-decay basidiomycete, in Europe mostly *Fomitiporia mediterranea* (Mugnai *et al*., 1999; Crous & Gams, 2000). Even if there are wood infections by some of these pathogens in the nursery (causing brown wood streaking and Petri decline, *sensu* Surico, 2010), in the vineyard wood tissue can get infected via pruning wounds by those vascular pathogens and the wood-decay basidiomycetes. These wood infections can remain latent throughout the vine’s life, or they can be associated with ‘grapevine leaf stripe disease’ (GLSD), a disease characterized by “tiger-striped” leaves and, less frequently, black measles on the berries, which is the most damaging disease within ECDs and among all GTDs in Europe, often simply called ‘esca’ (Larignon & Dubos, 1997; Mugnai *et al*., 1999; Graniti *et al*., 2000 ; Bertsh *et al*., 2013 ; Kandić *et al*., 2026). Wood-affected vines may also show sudden wilting of the canes or partial or total apoplectic symptoms. The development of foliar symptoms associated with wood infection shows a strong variability among cultivars and throughout the season, and the modality by which the wood pathogens cause GLSD is not yet fully clear.

Long-term field observations within single vineyards have shown that both symptoms incidence and severity fluctuate markedly from year to year (Marchi *et al*., 2006; Calzarano *&* Di Marco, 2007; Andreini *et al*., 2014; Calzarano *et al*., 2014; Ouadi *et al*., 2019; Moret *et al*., 2021; Del Frari *et al*., 2022) while vines with wood infections can remain asymptomatic for years or even for all their life (Calzarano & Di Marco, 2007; Romanazzi *et al*., 2009). Actually, the alternation of symptomatic and asymptomatic foliar expression from year to year within the same plant appears, following most authors, to be largely independent of the extent of internal wood symptoms (Calzarano *&* Di Marco, 2007; Romanazzi *et al*., 2009; Surico *et al*., 2010; Maher *et al*., 2012). However, vines that have even just occasionally expressed foliar symptoms show a significantly higher mortality rate compared with always asymptomatic plants (Guérin-Dubrana *et al*., 2013; Serra *et al*., 2021; Dewasme *et al*., 2022).

The erratic nature of GLSD symptoms suggests that abiotic factors play a major role in symptoms triggering. Among these factors, light exposure, soil physical and chemical properties, nutrient inputs through fertilisation, and water supply from rainfall and soil reserves have all been proposed as key modulators of symptoms expression (Mugnai *et al*., 1999; Calzarano *&* Di Marco, 2007; Calzarano *et al*., 2009, 2014). Among abiotic drivers, variations in water availability during the growing season have been consistently identified as a primary driver associated with the interannual alternation of symptomatic and asymptomatic years. Water availability consequent to rainfall patterns and leaf symptoms expression are strongly correlated; abundant precipitation, particularly during spring-early summer (May-July), is positively correlated with increased annual incidence of foliar symptoms (Surico *et al*., 2000a; Marchi *et al*., 2006; Calzarano *et al*., 2018). Experimental studies conducted under controlled conditions provide complementary evidence. Surico *et al*. (2010) reported a significant increase in foliar symptoms in 20-year-old potted vines, affected by GLSD, when subjected to excessive irrigation, whereas vines exposed to drought stress did not develop ‘esca leaf symptoms’ (i.e. GLSD) in a parallel experiment (Bortolami *et al*., 2021). These findings reinforce the central role of the water regime in the development of GLSD foliar symptoms.

More recent large-scale studies have further highlighted the importance of soil-water interactions. Using a statistical modeling approach applied to 493 vineyards planted with ten different cultivars across France between 2003 and 2022, Etienne *et al*. (2025) identified key ecoclimatic factors influencing annual GLSD incidence. Disease incidence increased significantly with higher soil wetness index values and evapotranspiration during the growing season, with the climatic conditions of the current year exerting a stronger influence than those of previous years. Similarly, an epidemiological study conducted on 19 vineyard plots across four Swiss wine-growing regions revealed a significant correlation between ‘esca’ incidence - particularly vine mortality - and soil water-holding capacity (Monod *et al*., 2025). These results are consistent with earlier findings suggesting that the ability of soils to retain water plays a crucial role in ‘esca’ development (Graniti *et al*., 2000; Van Niekerk *et al*., 2011; Calzarano *et al*., 2018; Calvo-Garrido *et al*., 2021).

Taken together, these findings provide strong evidence that water availability is a primary driver of GLSD foliar symptoms expression. We also know that soil properties, particularly physical properties, play a key role in regulating vine water status by modulating water retention and movement within the soil profile, thereby shaping its availability to the root system (Saxton *&* Rawls, 2006). Climate, particularly mean annual precipitation, subsequently influences soil chemical properties by altering pH, promoting nutrient leaching, and modifying redox conditions.

Despite all these evidences, knowledge regarding the specific role of soil chemical, physical and hydrological properties in GLSD incidence remains limited. An important premise to this understanding is the proven difference between the drivers of apoplectic strokes and GLSD (Magnin-Robert *et al*., 2016), a distinction that is not always specified when discussing the relationship between soil and ‘esca’. One of the few studies addressing this aspect (Corti *et al*., 2007) reported that soils associated with high ‘esca’ incidence were characterized by severe erosion leading to progressive nutrient depletion. In addition, soils with a high content of expandable clay (smectite group) induce summer soil cracking, causing root damage and desiccation, as well as poor drainage conditions during winter and spring. This study also reported that low phosphorus availability, low organic matter content, and excessive copper accumulation can increase the risk of ‘esca’ foliar symptoms development. Conversely, high nutrient availability has also been implicated in GLSD development. Nitrogen fertilization has been shown to increase disease incidence (Dell’Acqua *et al*., 2025), and similar effects have been reported for NPK fertilization and the excessive supply of micronutrients (Calzarano *et al*., 2023). Under humid climatic conditions, such as those in Swiss vineyards, Monod *et al*. (2025) found clear correlations between soil available water-holding capacity (AWC) and “esca disease” foliar and apoplectic symptoms.

Collectively, these findings could indicate that GLSD foliar symptom expression results from a complex interplay among climatic conditions, soil physicochemical properties, and vine physiology. Although rainfall is recognized as a key trigger of GLSD, the exact mechanisms it activates remains unexplained. Given the profound influence of rain on soil chemical, physical, and biological properties, it is worth to investigate wether soil-related factors may play a role in mediating symptom expression.

Investigating these relationships requires accounting for soil spatial variability within vineyards, as symptoms development often appears completely random. Apparent soil electrical conductivity (ECa), measured via electromagnetic induction (EMI), provides an integrative indicator of this variability, reflecting the combined influence of soil texture, soluble salts, water content, bulk density, porosity, soil depth, and organic matter (Corwin & Lesch, 2005). When combined with direct soil observations, ECa mapping can drive the selection of contrasting sampling locations and support the interpretation of soil heterogeneity (Priori *et al*., 2013). Several studies have demonstrated strong relationships between ECa and key edaphic properties, confirming its reliability for characterizing within-field spatial variability in agricultural systems, including vineyards (McLachlan *et al*., 2022; De Feudis *et al*., 2025). This approach provides a robust basis for selecting soil profiles in areas with contrasting histories of GLSD symptom expression.

To address this knowledge gap, the present study examines the associations between soil physicochemical properties and GLSD symptoms recurrence within a single vineyard monitored over five consecutive years. An EMI survey was used to delineate areas with contrasting apparent electrical conductivity (ECa), thereby identifying different soil types. These areas were overlaid with mapped high recurrence of symptomatic vines areas (HRSVAs) and low recurrence of symptomatic vines areas (LRSVAs) throughout the five-year monitoring period. Four soil profiles were described through trench excavation: two in HRSVAs and two in LRSVAs, each including sites with contrasting ECa values.

To the best of our knowledge, this is the first study to combine detailed pedological characterisation across multiple soil depths with comprehensive soil physicochemical analyses in relationship with GLSD symptom expression at a specific site. This study aims to clarify wether distinct soil properties are associated with varying GLSD symptom occurrences, with the ultimate aim of identifying how rainfall influence symptom expression. Complementary analyses of rhizosphere microbial communities are reported separately (Bigazzi *et al*., submitted).

## Materials and methods

### 1. Site description

The study was carried out in a ∼1.75 ha commercial vineyard of *Vitis vinifera* L. cv. Cabernet Sauvignon (Kober 5BB rootstock) located in the Bolgheri viticultural area (Figure 1) (Tuscany, Central Italy), a coastal wine-growing district officially recognised as a D.O.C. since 1983. The vineyard, established 25 years prior to the study, was trained to a spur-pruned cordon system with a spacing of 2.5 m between rows and 0.8 m between vines (5000 vines ha ¹). Canopy management included summer pruning and shoot trimming. The vineyard was managed without irrigation. Fertilisation consisted exclusively of on-farm compost produced from cattle manure composted for 10 months with straw, grape pomace, grape stems and shredded vine prunings. Interrows were managed alternately by soil tillage or by sowing a mixture of legumes, brassicas and grasses, which was incorporated into the soil in spring as green manure. The vineyard is non-irrigated and is located on level terrain within a system of Quaternary alluvial fan and terraces. The most prevalent soil type in this area is classified as *Endostagnic Luvisols* (IUSS Working Group WRB, 2022), as reported by the Tuscany regional soil map (Regione Toscana, 2017). These soils are distinguished by the process of clay illuviation, where clay particles move from the topsoil to deeper layers. In addition, the Endostagnic qualifier highlights the presence of reducing conditions due to temporary waterlogging between 50 and 100 cm. They have a neutral pH level and are often calcium carbonate free. Additionally, their texture varies, ranging from sandy loam to clay loam.

**Figure 1.**
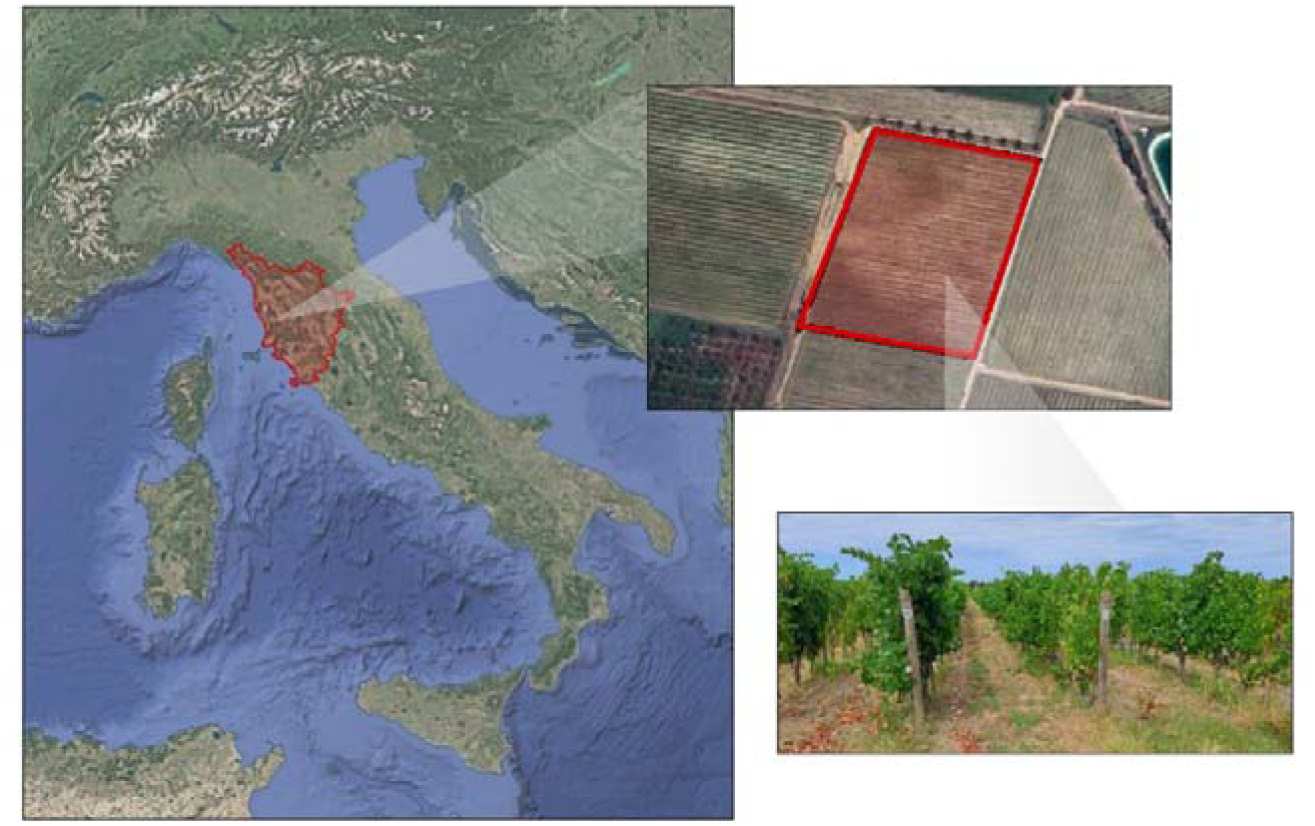
Vineyard selected as the study site, located in Bolgheri (Tuscany, Italy). The red box highlights the vineyard plot that was monitored for foliar symptoms of ECDs.

The area is characterized by a Mediterranean climate with mild, wet winters and warm, dry summers, with an average annual temperature of 14.4–15.5 °C and annual rainfall of approximately 500–600 mm, less than 200 mm of which occurs between April and August (https://www.sir.toscana.it).

The vineyard is equipped with a weather station that records rainfall data. Figure 2 shows cumulative rainfall (mm) during May–July and annual GLSD foliar symptom incidence from 2020 to 2024, providing meteorological and disease context for the study site. This period was selected because previous studies have associated rainfall during late spring and early summer with GLSD foliar symptom expression (Marchi *et al*., 2006; Calzarano *et al*., 2018). As illustrated in the graph, the annual incidence of foliar symptoms follows the rainfall trend observed during these months.

**Figure 2.**
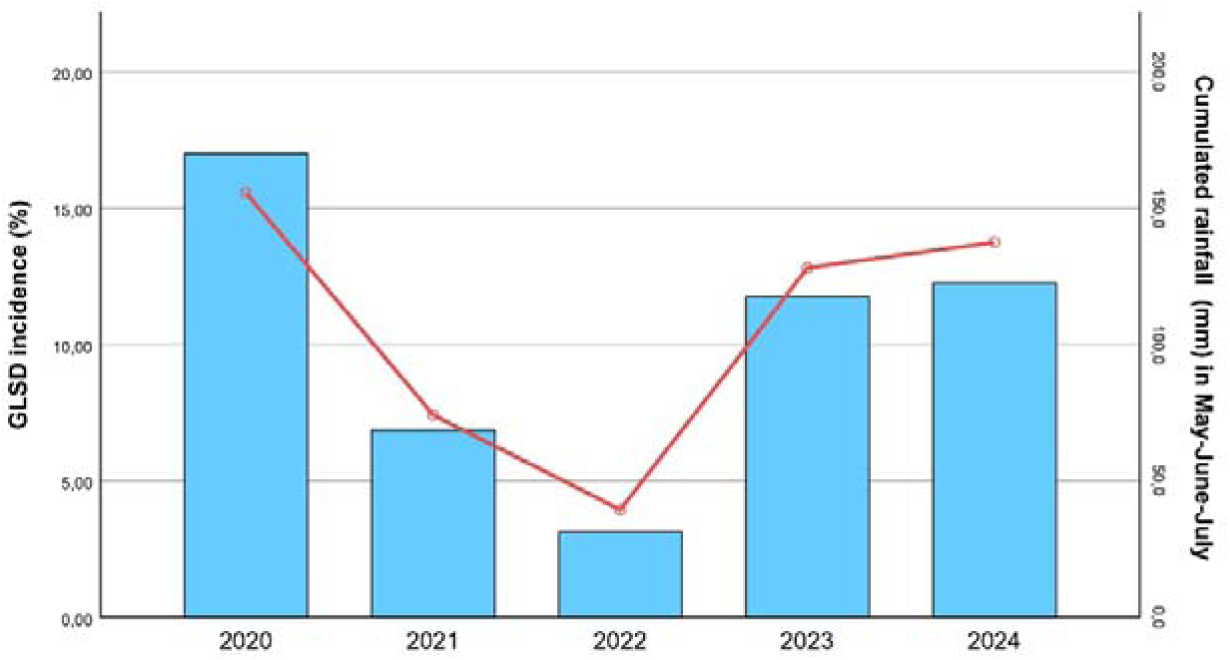
Annual incidence of Grapevine Leaf Stripe Disease (GLSD) foliar symptoms (red line, left y-axis) and cumulative rainfall recorded during May, June, and July (blue bars, right y-axis) in the Bolgheri vineyard over the 2020–2024 period.

### 2. Grapevine leaf stripe disease symptoms evaluation and mapping

Grapevine Leaf Stripe Disease (GLSD) symptoms were assessed through field surveys conducted annually from 2020 to 2024 (Figure 3). Monitoring covered 71 rows within an area of approximately 1.75 ha, comprising approximately 10,000 standing vines at the first survey. Surveys were conducted in September, when foliar symptoms are generally well developed and readily detectable. Vines exhibiting typical tiger-striped leaves were classified as symptomatic, irrespective of whether shoot wilting was also present. Shoot wilting, including the few cases classified as apoplexy (defined here as >60% canopy wilting), was recorded only in association with tiger-striped leaves and was not used as an independent criterion for symptom classification. Thus, all vines classified as symptomatic exhibited tiger-stripe symptoms, although some also showed shoot wilting or partial canopy collapse. Some vines died during the five-year monitoring period. Vines that died following the expression of GLSD foliar symptoms recorded during monitoring were included among diseased vines. Each vine was georeferenced and mapped as point shapefile using QGIS software, and a cumulative disease incidence score was assigned on a scale from 0 to 5, corresponding to the number of years in which GLSD symptoms were observed during the monitoring period.

**Figure 3.**
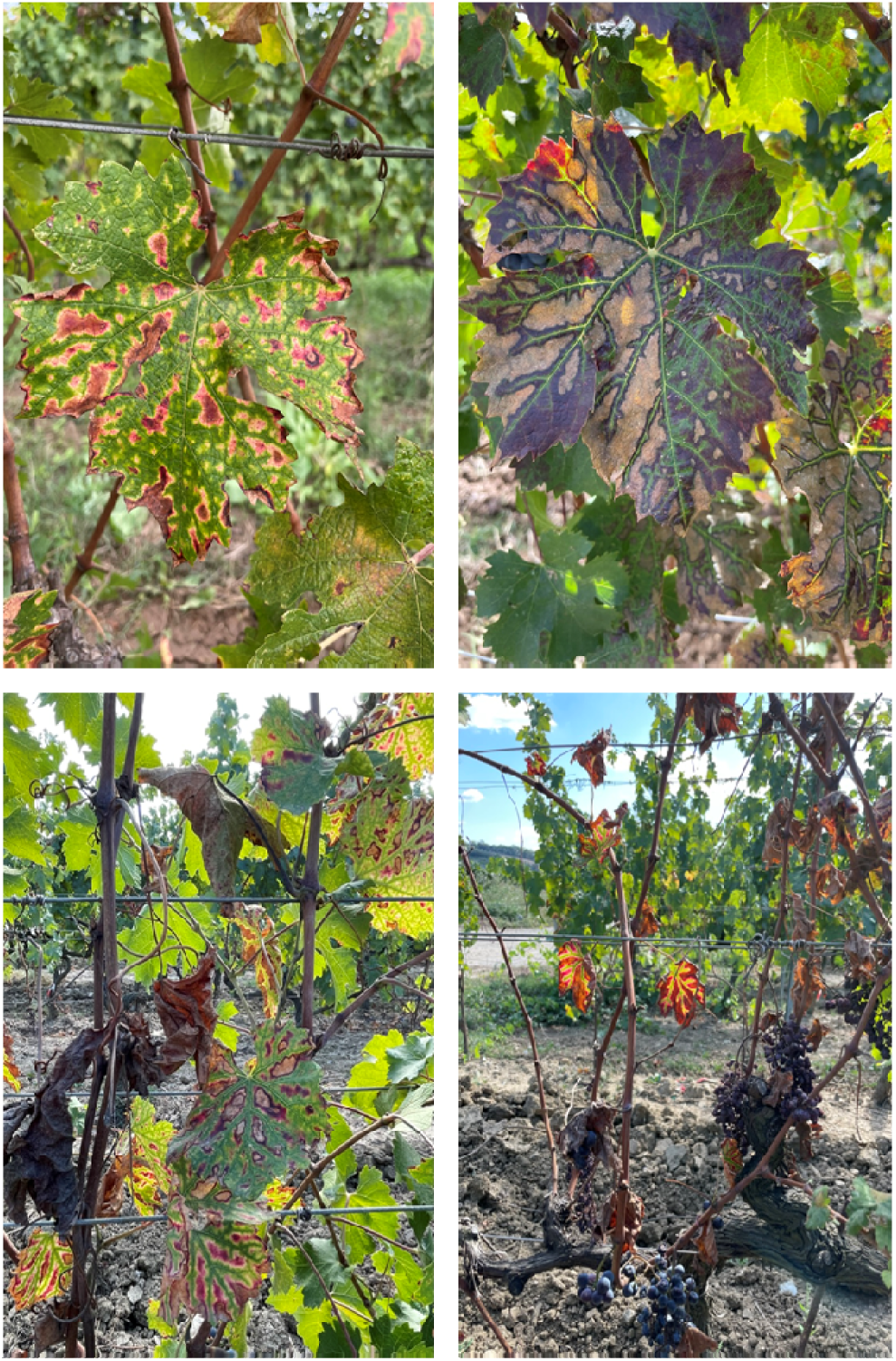
Top: typical foliar “tiger stripe” symptoms of grapevine leaf stripe disease. Bottom left: shoot wilting with tiger-stripes leaves. Bottom right: apoplexy symptoms with tiger striped leaves.

Maps of GLSD symptoms recurrence within 5 years were carried out using Ordinary Kriging on dummy variables, to simulate the method of Indicator Kriging (Journel, 1983). The cumulative disease incidence of the monitored vines was transformed into dummy variables (binary, 0 for absence, 1 for presence) indicating: i) at least one incidence in 5 years; ii) at least 3 incidences in 5 years. These dummy binary variables were interpolated using ordinary block kriging, with block size of 5×5 m. The semivariograms were modelled using a short range (12 m) and a short lag (1 m) because of the high spatial variability of the symptoms. The models used were both exponential.

One map was set up to show the probability of recurrence of GLSD symptoms appearing at least once within a five years period, whereas the other was set up to display the probability of encountering the same symptoms on at least three of the five years.

The interpolation with ordinary block kriging was carried out by SAGA Gis software (version 9.0), whereas the map editing and layout for final figures was carried out by QGis software (version 3.44).

### 3. EMI proximal sensing and sampling design

Soil spatial variability was mapped using the Mini Explorer, an electromagnetic induction sensor produced by GF Instruments (Brno, Czech Republic). It consists of a system of transmitting (Tx) coil, which operates at 30 kHz, and three receiving parallel coils (Rx), spaced 0.32, 0.71 and 1.18 m from the Tx. This instrument configuration investigates the apparent electrical conductivity (ECa) of the soil at 3 depths: ECa_1_ around 0-50 cm, ECa_2_ around 0-100 cm, and ECa_3_ around 0-150 cm. Like the other commercial EMI sensors, the GF Mini-Explorer is widely used in agriculture to investigate and describe spatial variations in the root zone volume of soils in vineyards and other arboreal crops (Pace *et al*., 2024; Lavaud *et al*., 2025). The sensor is equipped with a GNSS system to georeference the points detected (1 measurement per second), and then to obtain maps of ECa_1_, ECa_2_, and ECa_3_ with a resolution of 1 m, interpolating the measurement points by ordinary kriging, after semivariogram modelling.

Based on soil ECa measurements, areas of relatively high and low conductivity were delineated within the vineyard. Spatial integration of cumulative GLSD incidence maps and soil ECa maps allowed the identification of vineyard sectors characterized by contrasting combinations of disease incidence and soil conductivity. This sampling strategy enabled the comparison of soils with contrasting ECa under similar levels of cumulative disease incidence, as well as soils with similar conductivity under contrasting disease pressure. This approach was adopted to evaluate whether soils characterized by different ECa conditions—reflecting potential differences in soil physical and hydrological features—share common soil properties in areas of high cumulative disease incidence that differentiate them from areas with low cumulative disease incidence.

Four areas within the vineyard were selected according to two criteria: cumulative disease incidence (high vs. low) and soil electrical conductivity (high vs. low), resulting in four contrasting combinations: (i) high incidence/low conductivity, (ii) high incidence/high conductivity, (iii) low incidence/low conductivity, and (iv) low incidence/high conductivity. Each areas is represented by a benchmark soil profile, excavated about 40 cm next to the vine row, as following: P1, low symptom recurrence and medium ECa; P2, low symptom recurrence and high ECa; P3, high symptom recurrence and medium ECa; and P4, high symptom recurrence and high ECa. Medium and high ECa refer to relative conductivity levels at the selected sites within the vineyard, rather than sharply contrasting or discrete soil classes.

To calculate the ECa values of the maps around each sampling location, a circular buffer with a diameter of 5 m was created around each profile location in SAGA GIS. The tool “Extract Grid Values from Shapefile” was used to extract the mean raster values within each buffer from the ECa maps (ECa1, ECa2 and ECa3) and the cumulative GLSD incidence map. This procedure provided a local mean for each mapped variable based on the raster cells within the buffer, rather than a value from a single pixel. The locations corresponding to the absolute minimum ECa values were excluded from profile selection because they occurred at the ends of vineyard rows. Sampling locations further inside the vineyard were therefore selected, resulting in profiles representing relatively lower (medium) and higher ECa conditions, rather than the absolute low and high extremes of the vineyard.

To verify the statistical correlation between cumulative GLSD incidence over the 5 years and the ECa maps, we perfomed and ANOVA test, using the number of years with GLSD symptoms as class (0 to 5) and ECa_1_, ECa_2_, and ECa_3_ as numerical variables.

### 4. Soil profiles description and analysis

At each selected location, a soil trench was dug to a depth of approximately 1.2 m adjacent to the vine row, on 8 November 2024, extending along a section of the row encompassing three grapevine. Soil profiles were described in the field, and horizons were identified according to the World Reference Base for Soil Resources (IUSS Working Group WRB, 2022). During the pedological description, particular attention was paid to pedogenetic features related to waterlogging, namely redoximorphic mottles, as well as Fe-Mn concentrations and nodules. In addition, a description of the grapevine root system was also done, to highlight eventual rooting limitations or particular roots concentration in specific soil horizons. The volumetric fraction (in %) of the coarse fragments were estimated in field for each horizon. For each soil horizon, three soil samples were collected along the trench wall, one adjacent to each of the three grapevines. The samples were subsequently combined to form a single composite sample representative of each horizon. Undisturbed soil samples for bulk density (BD) were collected for each horizon until 1 m of depth, by steel rings of 100 cm^3^. The laboratory analysis of soil samples includes: particle size distribution, pH, electrical conductivity (EC), total and active lime, organic carbon (OC), total nitrogen (TN), cation exchange capacity (CEC), exchangeable calcium (Ca), magnesium (Mg), potassium (K), and sodium (Na), as well as available phosphorus (P), iron (Fe), manganese (Mn), and soluble boron (B). All soil analyses were performed according to official Italian analytical methods for soil chemical and physical analysis (Ministerial decree n. 185, 13/9/1999). In particular: particle size distribution was carried out by hydrometer method; pH and EC in water solution (1:2.5 and 1:5, respectively); active lime was determined by ammonium oxalate extraction and permanganate back-titration; total and organic carbon, as well as total nitrogen by CN analyzer; CEC was measured by ammonium saturation using 1 M ammonium acetate at pH 7.0; exchangeable Ca, Mg, K and Na were extracted with ammonium acetate and quantified spectrometrically; available phosphorus was determined by Olsen method. Available Fe and Mn were extracted with ammonium bicarbonate–DTPA and DTPA–CaCl□–triethanolamine buffered at pH 7.3, respectively, and measured by atomic absorption spectrometry, whereas available B was extracted by CaCl and determined by ICP-AES. The undisturbed soil samples were dried to 105 °C for 24 h and weighted, to determine the bulk density of each horizon. Available water capacity (AWC, in mm of water in 1 m of soil), and saturated permeability (K_sat_) are calculated for each soil horizon using the pedofunctions of Saxton and Rawls (2006), implemented in the software SPAW (Soil-Plant-Air-Water, USDA-NRCS, https://www.nrcs.usda.gov/resources/tech-tools/spaw-version-602). In particular, the pedofunctions calculated the water content at Field Capacity (θ_33_) and Wilting Point (θ_1500_) as :

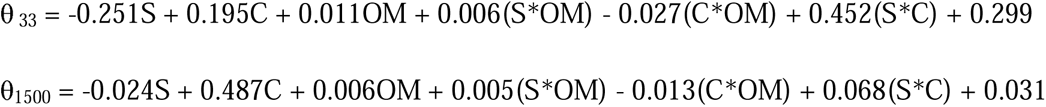

where S and C are the sand and clay fractions (%), OM is the organic matter (g 100 g^-1^). Soil organic matter content (OM) was estimated by multiplying soil organic carbon content by the conventional van Bemmelen factor of 1.724, assuming a carbon content of 58% in soil organic matter (Nelson *&* Sommers, 1996). Both θ_33_ and θ_1500_ were corrected for bulk density. AWC was calculated as the difference between θ_33_ and θ_1500,_ and then corrected for the coarse fragments (Saxton *&* Rwals, 2006). The profile AWC was calculated until a depth of 1 m as average, weighted for horizon thickness. The saturated hydraulic conductivity (Ksat) of the entire soil profile was estimated using the thickness-weighted harmonic mean of the individual horizons, as described by Hillel (1998).

## Results

### 1. Distribution of cumulative symptoms distribution

The distribution of annual GLSD symptoms was very heterogeneous, with a symptomatic vine next to asymptomatic vines (Figure 4). On the other hand, areas with higher recurrence were clearly observable, after the spatialization of the cumulative incidence data on yearly symptomatic observations by indicator kriging (Figure 5). It is interesting to note that the symptoms recurrence map of one symptomatic year within 5 years of observation (Figure 5a) showed heterogeneous pattern, with a general major recurrence of the symptom in the southern part of the vineyard. On the other hand, the map showing the recurrence of GLSD symptoms appearing in at least three of the 5 monitored years (Figure 5b), showed a clearer spatial pattern. Distinct areas characterized by higher GLSD symptom recurrence were observed, particularly in the southern and eastern portions of the vineyard. An area with lower symptom recurrence was detected in the central sector of the vineyard and in the north-western portion of the parcel.

**Figure 4.**
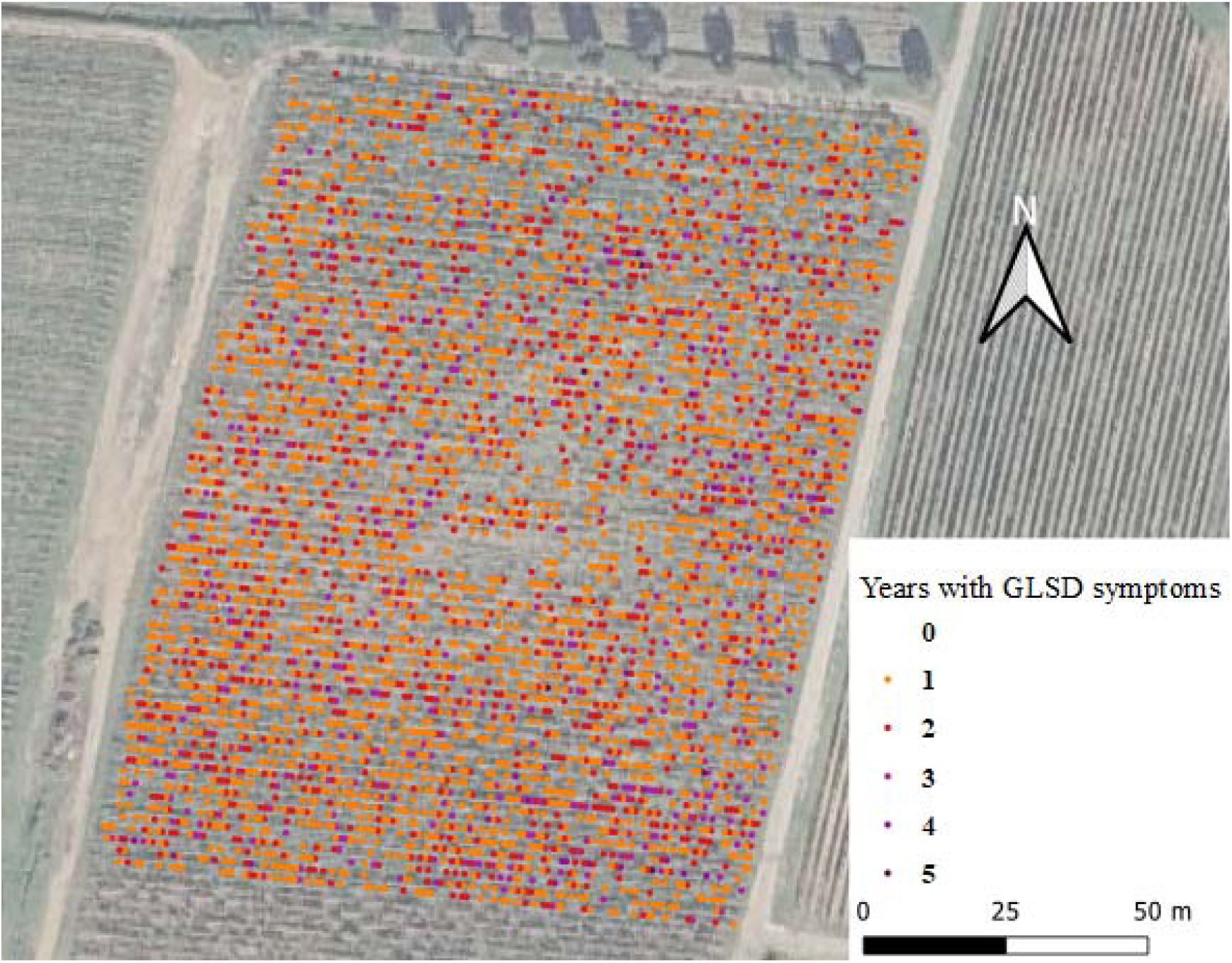
Map showing all the grapevines monitored and how many years each vine has shown symptoms of GLSD. Vines that have never shown symptoms are marked with a transparent symbol. The values of this figure are real, not interpolated.

**Figure 5.**
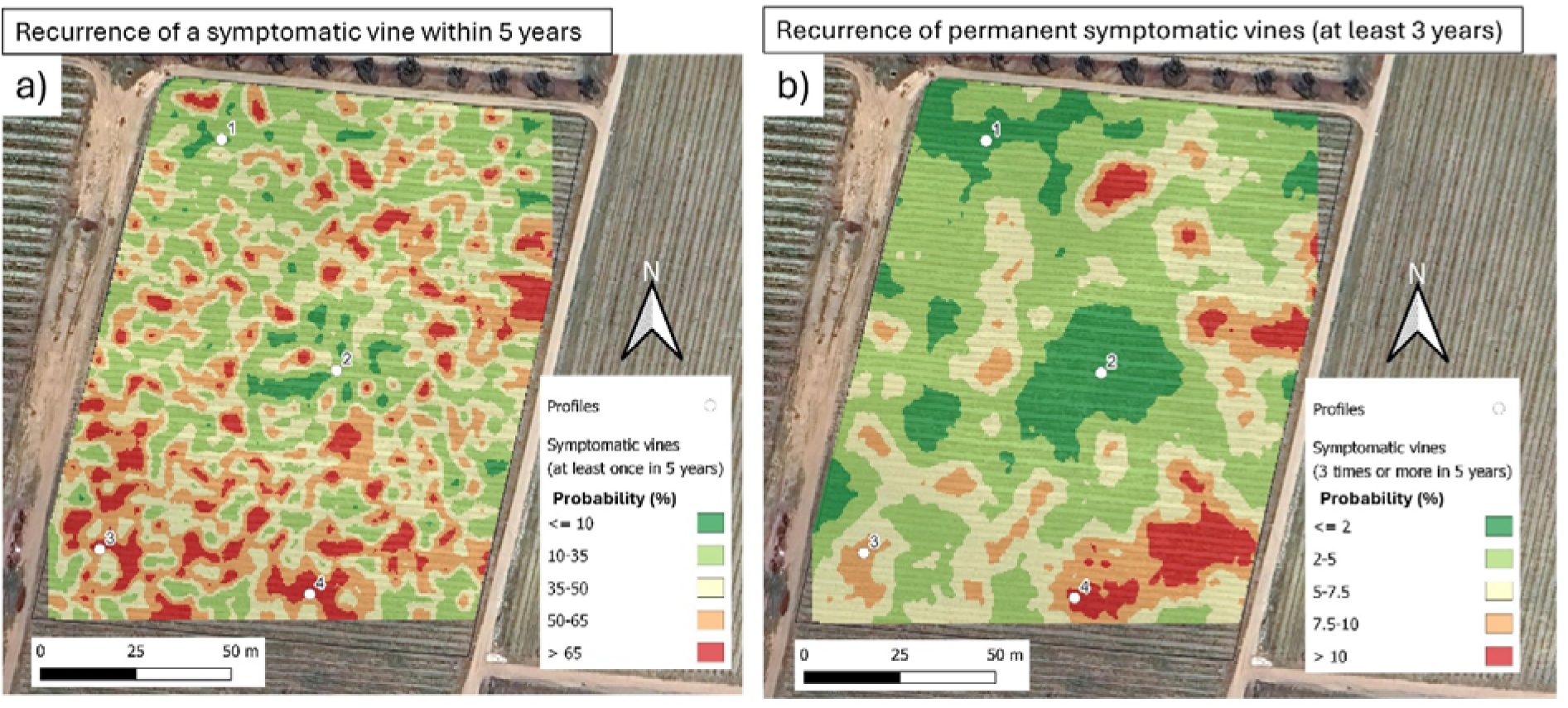
Maps showing the probability of GLSD symptom recurrence over the five-year monitoring period within the vineyard parcel, obtained by ordinary block kriging of binary indicator variables. The map on the left (a) shows the probability of encountering vines that exhibited GLSD symptoms at least once during the five-year monitoring period, whereas the map on the right (b) shows the probability of encountering vines that exhibited GLSD symptoms in at least three of the five monitoring years.

### 2. ECa maps and soil profiles location

The EMI proximal survey data, interpolated by ordinary kriging, produced the maps of apparent electrical conductivity (ECa) at three depths: ECa_1_ 0-50, ECa_2_ 0-100, and ECa_3_ 0-150 cm (Figure 6). The maps showed a similar pattern, characterized by a central part of the vineyard showing high ECa values, from 35-40 mS·m^-1^ (ECa_1_) to 80-110 mS·m^-1^ (ECa_3_). Such high ECa values are typical of deep, moist, clayey soils (Pedrera-Parrilla *et al*., 2016; Pace *et al*., 2024), and are likely not driven by salinity, which generally results in readings higher than 200 mS·m^-1^ (Doolittle *&* Brevik, 2014). The lack of salinity was confirmed by the soil analysis, reported in Table 1.

**Figure 6.**
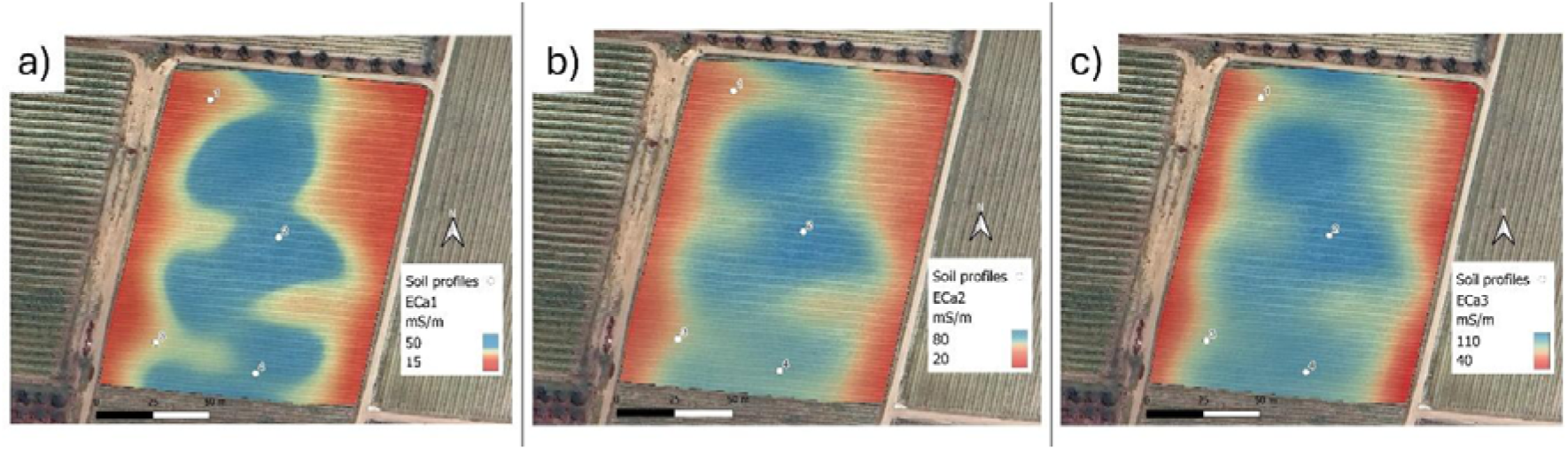
Maps of apparent electrical conductivity (ECa) measured by GF Mini-Explorer sensor. The maps interpolate points of measurement at 3 depths (ECa1 at 0-50 cm; ECa2 at 0-100 cm, ECa3 at 0-150 cm) by ordinary kriging, and shows the position of the fours soil profiles, P1, P2, P3 and P 4.

**Table 1.** Mean ECa values and mean mapped cumulative GLSD incidence within a 5-m-diameter buffer centred on each soil-profile location. ECa1, ECa2 and ECa3 correspond to approximate investigation depths of 0–50, 0–100 and 0– 150 cm, respectively. GLSD incidence (%) represents the mean mapped percentage of vines showing symptoms at least once during the five-year monitoring period (2020–2024).

| Profile | Area | Relative ECa | ECa1<br>(mS m<sup>-1</sup>) | ECa2<br>(mS m<sup>-1</sup>) | ECa3<br>(mS m<sup>-1</sup>) | GLSD recurrence<br>(%) |
| --- | --- | --- | --- | --- | --- | --- |
| P1 | LRSVA | Medium | 29.3 | 52.1 | 74.1 | 23 |
| P2 | LRSVA | High | 44.1 | 81.3 | 115.0 | 20 |
| P3 | HRSVA | Medium | 30.3 | 51.4 | 74.6 | 57 |
| P4 | HRSVA | High | 37.7 | 66.8 | 99.2 | 67 |

The analysis of variance of the ECa values and GLSD symptoms recurrence showed no significant correlations, and this is clearly observable comparing the two maps of Figure 5. Therefore, the ECa maps cannot be used as predictive variables to assess the GLSD recurrence. On the other hand, ECa maps were used to locate the soil profiles and to determine the soil spatial variability of the vineyard.

The eastern and western borders of the vineyard showed lower ECa values, ranging from approximately 15 to 30–40 mS m ¹, indicating soils with slightly lower clay content and moisture, as well as higher gravel and sand content, as confirmed by the description and soil analyses of P1 and P3 (Table 2). The absolute minimum ECa values occurred near the ends of vineyard rows; these marginal locations were therefore excluded, and sampling locations further inside the vineyard were selected (Figure 4). The soil profiles were located according to the variability of ECa and the recurrence of symptomatic vines (Figure 7), as follows:

**Figure 7.**
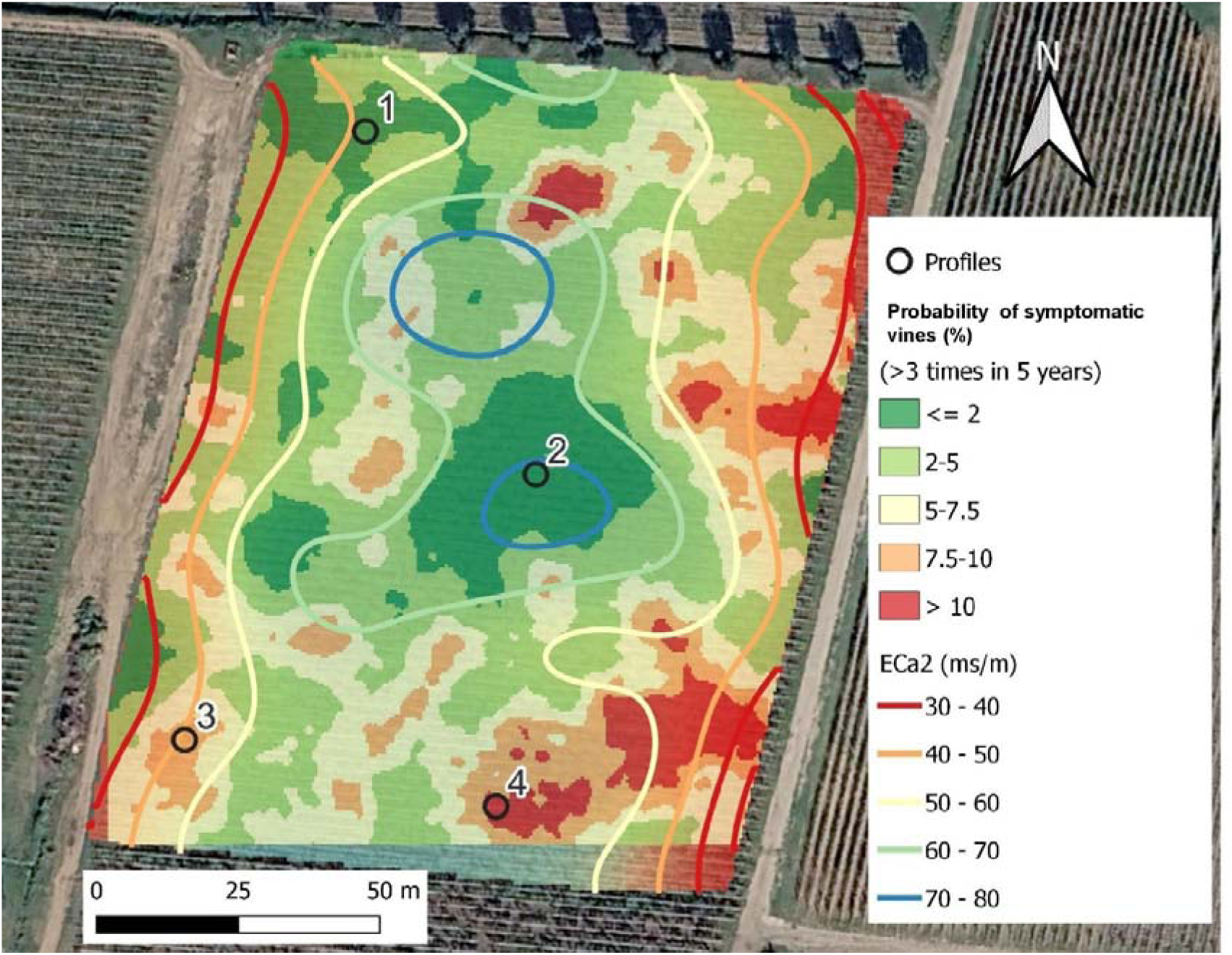
Identification of soil profiles based on the probability of GLSD symptom recurrence and apparent soil electrical conductivity (ECa).

**Table 2.**
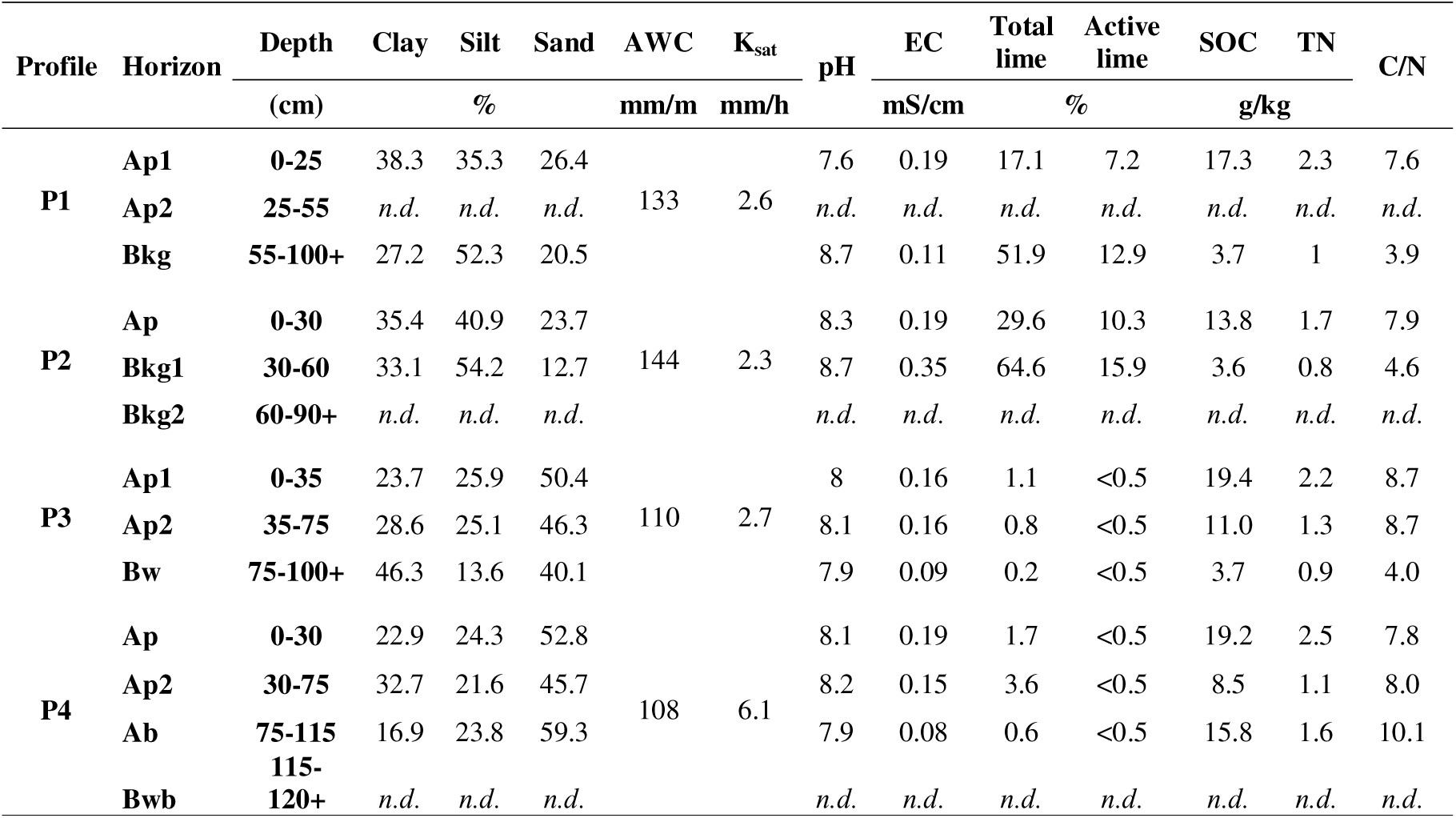
Physical, hydrological, and chemical features of the soil profiles.

P1: low recurrence of symptomatic vines (LRSVAs; ≤5% probability of encountering vines that exhibited GLSD symptoms in at least three of the five monitoring years), medium ECa (30, 50 and 73 mS m□¹ for ECa1, ECa2 and ECa3, respectively).

P2: low recurrence of symptomatic vines (LRSVAs; ≤5% probability of encountering vines that exhibited GLSD symptoms in at least three of the five monitoring years), high ECa (45, 82 and 116 mS m□¹ for ECa1, ECa2 and ECa3, respectively).

P3: high recurrence of symptomatic vines (HRSVAs; ≥7.5% probability of encountering vines that exhibited GLSD symptoms in at least three of the five monitoring years), medium ECa (30, 52 and 75 mS m□¹ for ECa1, ECa2 and ECa3, respectively).

P4: high recurrence of symptomatic vines (HRSVAs; ≥7.5% probability of encountering vines that exhibited GLSD symptoms in at least three of the five monitoring years), high ECa (37, 67 and 100 mS m□¹ for ECa1, ECa2 and ECa3, respectively).

The mean ECa values and mapped cumulative GLSD incidence associated with each selected profile are reported in Table 1. Across the vineyard, ECa ranged from 21 to 46 mS m□¹ for ECa1, 37 to 84 mS m□¹ for ECa2 and 50 to 125 mS m□¹ for ECa3, while cumulative GLSD incidence ranged from 4% to 80%.

### 3. Soil characteristics

The soils of the profiles P1 and P2, situated in LRSVAs, showed high similarity and differed from the other soil profiles P3 and P4, situated in HRSVAs. In particular, both P1 and P2 showed carbonatation process in subsoil horizons (Bkg horizons) and very high amount of calcium carbonate in depth, more than 50% of total lime and more than 12% of active lime (Table 2). Both the profiles showed clay loamy or silty-clay loamy texture and slow internal drainage, with temporary waterlogging under 30-55 cm, testified by high frequency of redoximorphic mottles (Figure 8). During the profile digging, a temporary water-table was found at 80 cm in P2 profile. The calculated AWC of P1 and P2 was higher (133 and 144 mm·m^-1^) than that in P3 and P4 (110 and 108 mm·m^-1^). The estimated soil K_sat_ was very similar for P1, P2, P3 (from 2.3 to 2.7 mm·h^-1^), defined as moderately slow by USDA-NRCS (2017), whereas P4 showed a value of moderately high class (6.1 mm·h^-1^).

**Figure 8.**
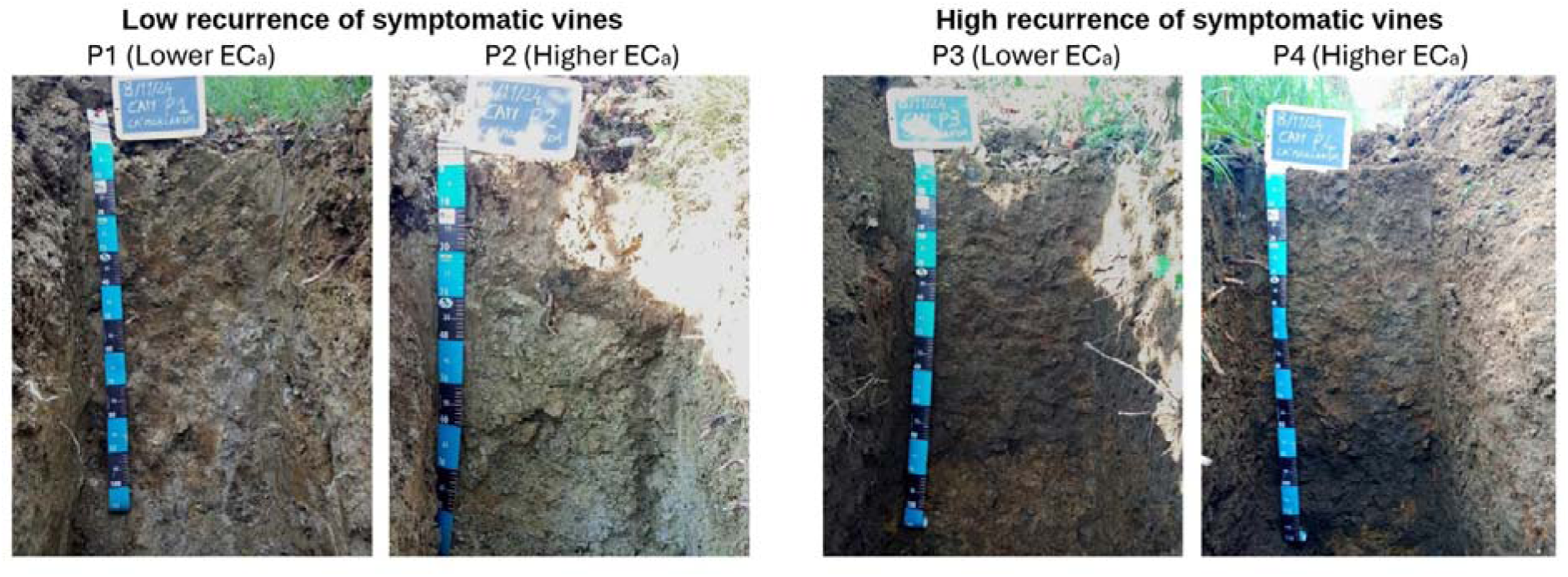
Soil profiles in areas with low recurrence of symptomatic vines (LRSVAs, left) and high recurrence of symptomatic vines (HRSVAs, right).

The pH was sub-alkaline in the topsoil (A horizon) and alkaline in depth (Bkg horizons). This alkalinity was also confirmed by moderately high amount of exchangeable Na^+^ in the B horizons of P2. Salinity was low, although Bkg horizon in P2 showed slightly higher values (0.35 dS/m) than the other horizons. Soil organic carbon (SOC) was moderately low in A horizons (0-25 or 0-30 cm), but very low (< 5 g kg□¹) in the subsoil, B horizons. C/N ratio was also very low in B horizons. According to international WRB classification (IUSS Working Group WRB, 2022), P1 was classified as *Stagnic Calcisol (Loamic, Hypercalcic)*, whereas P2, because of its sodicity, was classified as *Stagnic Calcisol (Loamic, Hypercalcic, Sodic)*. Both the profiles showed clear limitations for grapevine growth and roots development, mainly due to the oxygen scarcity in depth, alkaline pH and high active lime.

The soils of the profiles P3 and P4, situated in HRSVAs, showed very different hydrological and chemical features than P1 and P2. The texture was variable from sandy loam to sandy clay loam, and clayey in the Bw horizon (90-100 cm) of P3 (Table 2). In these profiles, total lime was very low (< 4%, often < 1%) and active lime was under detection threshold (< 0.5%). The SOC is slightly higher than in P1 and P2, both in the topsoil (A horizons) and in the subsoil (B horizons). In particular, P4 showed a buried Ab horizon between 75 and 115 cm, with 15.7 g kg ¹of SOC, which was a high amount for this depth. This buried horizon could be probably due to the land levelling during the past vineyard planting. P3 did not show a clear Ab (buried) horizon, but a subsoil Ap2 (ploughed) horizon that may have originated by a deep tillage and mixing between original A and B horizons. Probably, during the land levelling for vineyard planting, P1 and P2 were the result of soil truncation, whereas P3 and P4 could be the result of the land filling of original depressions. In any case, P3 and P4 soils showed higher fertility than P1 and P2, not only in terms of total organic carbon and nitrogen, but also in terms of rooting explorability. If in P1 and P2, the grapevine roots were more concentrated in the first two horizons (0–60 cm), in P3 and P4, they were well distributed up to the maximum profile depth of 120-130 cm. According to international WRB classification (IUSS Working Group WRB, 2022), both P3 and P4 were classified as *Cambic Phaeozems (Loamic, Relocatic)*.

A clear difference between P3-P4 and the soils with low recurrence of symptoms (P1 and P2) was also the higher availability of Mn, in particular in the deeper horizons. Taking into account all the analysed horizons, the weighted average of available Mn was 4.5-4.8 mg kg ¹in P1 and P2, and 14.7-14.8 mg kg ¹in P3 and P4 (Table 3).

**Table 3.**
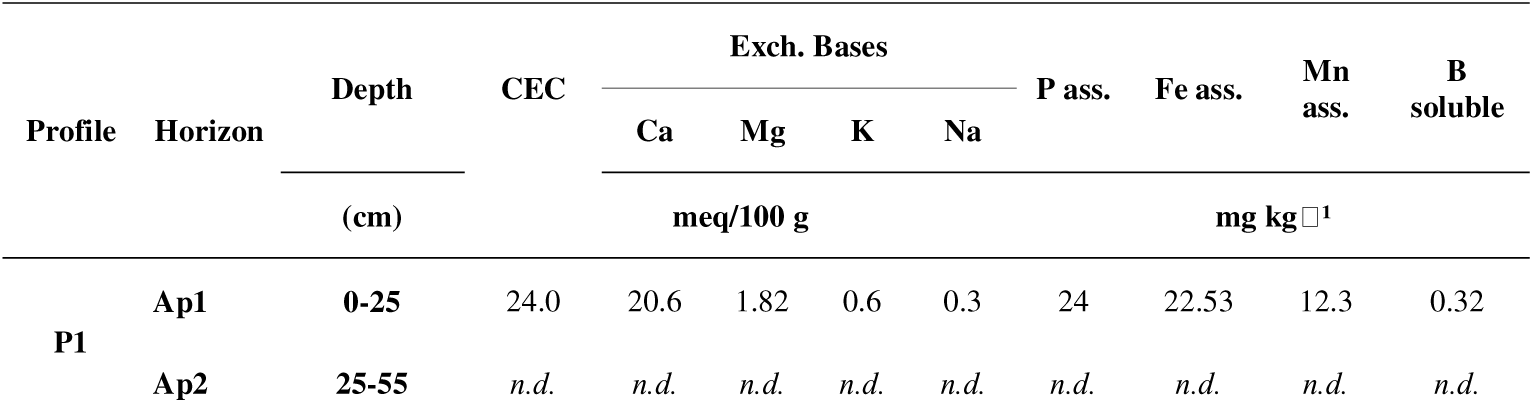
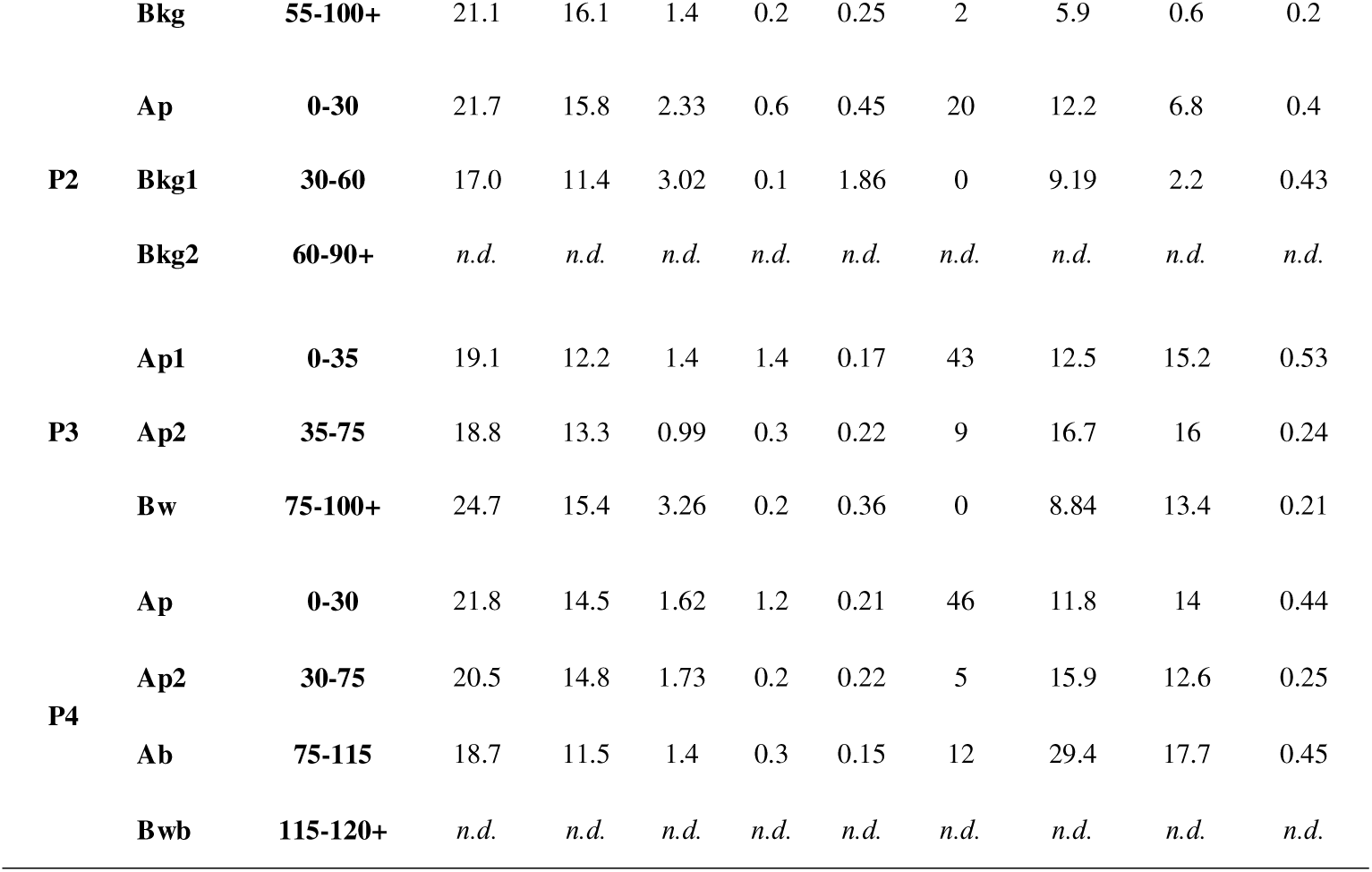
Cation Exchange Capacity, exchangeable bases and available micronutrients of the soil horizons.

## Discussion

The research here carried out supports some important findings. The non-uniform spatial distribution of GLSD symptoms within a single vineyard showed that recurrent symptoms can be associated with localized soil conditions within the vineyard parcel. Because of these different soil conditions, rainy springs can influence nutrients availability, and in particular the data showed that the differences detected in the soil of areas with high symptoms recurrence are strongly related to manganese availability, a nutrient that was shown to influence wood enzymatic degradation thanks to its role in manganese peroxidase.

Previous studies have shown a random spatial distribution of GLSD (or ‘esca’-related) foliar symptoms (Redondo *et al*., 2001; Sofia *et al*., 2006), whether other studies have detected some symptom aggregation both along or across rows (Surico *et al*., 2000; Stefanini *et al*., 2000; Pollastro *et al*., 2000; Bonneau *et al*., 2017). The spatial clustering has been primarly discussed in relationship to infection spread via contaminated pruning scissors (Surico *et al*., 2000; Stefanini *et al*., 2000).

Another peculiar aspect of GLSD is its foliar symptoms fluctuation over the years in the same vines. Since asymptomatic and symptomatic plants often host highly similar wood mycobiomes (Hofstetter *et al*., 2012; Bruez *et al*., 2014; Del Frari *et al*., 2019), the occurrence of foliar symptoms in a given season can not only depend on pathogen presence. This suggests a major role of abiotic factors (Graniti *et al*., 2000) in regulating pathogens activity. For example Surico *et al*. (2000a) observed a lower disease incidence on steeper slopes and higher in flatter areas, suggesting a topographic influence, probably linked to soil fertility and water availability, which are generally higher in flat areas. This is consistent with the wellknown relationship between rainfall and disease expression (Marchi *et al*., 2006; Monod *et al*., 2025).

Our five-year monitoring also showed a qualitative correspondence between rainfall and annual GLSD incidence, with rainy springs coinciding with greater symptom expression.

A more recent study conducted in French vineyards by Li *et al*. (2017) also supported the variability of spatial patterns in GLSD symptoms distribution. While some vineyards exhibited structured distribution of symptomatic plants, others showed random patterns. Importantly, when spatial structuring was observed, it was often associated with heterogeneity in soil composition and structure (Li *et al*., 2017). This interpretation is consistent with previous findings indicating that the incidence of foliar symptoms can be influenced by soil chemical properties and foliar fertilization practices (Calzarano *et al*., 2009). Abiotic factors, including nutrient availability, water supply, and temperature have in the past been been identified as possible drivers of yearly fluctuation in symptoms (Graniti *et al*., 2000; Corti *et al*., 2007; Fischer *&* Peighami-Ashnaei, 2019).

Building on this evidence, the results obtained in the present study further support the role of soil heterogeneity in shaping the spatial distribution of GLSD symptoms. In our work, proximal soil sensing using electromagnetic induction (EMI) revealed a marked variability in soil electrical conductivity (ECa) within the monitored parcel. Specifically, two distinct zones were identified, characterized by contrasting ECa values: a higher ECa zone in the central part of the vineyard and a lower ECa zone in the eastern and western parts. To investigate this further, two soil profiles were analysed within each of the two areas: one characterised by a low recurrence of symptomatic vines (LRSVAs) and the other by a high recurrence of symptomatic vines (HRSVAs). These profiles revealed substantial pedological differences between the areas with low and high recurrence of GLSD symptoms.

Regarding soil physical-hydrological parameters, profiles located in LRSVAs (P1 and P2) showed finer texture (clay loam to silty clay loam), higher available water capacity (AWC), but also slow drainage and strong waterlogging around 50-60 cm. The relativly higher AWC in LRSVAs contrasts with the positive correlation between GLSD and soil water holding capacity (SWHC) reported by Monod *et al*. (2025). However, their study was carried out in soils developed on morain deposits in Western Switzerland, characterized by higher coarse fragments and coarser texture than the soils studied in our research. Although the authors reported SWHC values similar to those of the present study, they calculated the water availability until 150 cm, instead of 100 cm, as in our data. SWHC values lower than 100 mm are considered low and generally can reduce the vine vigour by limiting plant water uptake, thereby potentially reducing the development of the GLSD symptoms. In our case, AWC varied from 108-110 mm/m in areas with higher GLSD recurrence (P3 and P4), and 138-144 mm/m in areas with lower GLSD recurrence (P1 and P2). On the other hand, rooting depth of P3 and P4 is deeper than 100 cm, making the water availability for plants higher than that measured. In this case, water availability did not seem to discriminate the areas with higher and lower GLSD recurrence. This is most probably the reason why in this specific study the AWC is not correlated with GLSD recurrence.

Furthermore our study indicates that higher AWC corresponds to increased waterlogging and greater difficulty for the vine to estabilish deep roots. Prolonged water saturation is known to reduce soil oxygen availability thereby restricting root deepening (Parent *et al*., 2008; Zhang *et al*., 2025). These internal conditions are physiologically comparable to those occurring under drought stress (Chen *et al*., 2023), an aspect that is particularly relevant in light of the findings of Bortolami *et al*. (2021), who observed that grapevines exposed to drought stress did not develop foliar symptoms, possibly because such water deficit limits the translocation of toxic fungal metabolites. More recently, Gastou *et al*. (2026), working on examining the same grapevines as Bortolami *et al*. (2021), demonstrated that this inverse relationship between drought conditions and GLSD symptoms expression was not associated with a reduction in grapevine wood pathogen abundance. These findings therefore support the hypothesis that drought limits the upward translocation of fungal toxic compounds and/or wood degradation by-products, which are believed to play a role in GLSD symptoms development (Mugnai *et al*., 1999; Andolfi *et al*., 2011), rather than affecting pathogen abundance itself.

In the present vineyard, the evidence of temporary waterlogging in P1 and P2 suggests that excess soil water may limit root exploration at depth, potentially reducing grapevine vigour and increasing susceptibility to summer drought through restricted rooting depth. Within this framework, the waterlogging conditions observed in LRSVAs (P1 and P2) can be coherently interpreted as functionally analogous to drought: by limiting root water uptake and reducing plant hydraulic conductivity (Chen *et al*., 2023), they may also restrict the upward movement of fungal metabolites or toward the canopy, thereby limiting foliar symptom expression despite pathogen presence. It is important to emphasise that pathogen presence alone does not necessarily lead to foliar symptom expression, as esca complex associated fungi have been detected also in asymptomatic vines (Hofstetter *et al*., 2012; Bruez *et al*., 2014).

Regarding soil chemical properties, a striking difference emerged in the profiles of LRSVAs (P1 and P2) which exhibited a high content of total and active lime, capable of nutritional stress of the plants (e.g. Fe and P nutrition, chlorosis). Conversly, HRSVAs were characterized by higher overall fertility (greater nitrogen and organic matter content) and greater root explorability. Dell’Acqua *et al*. (2025) reported that increasing nitrogen fertilization levels led to a higher incidence of GLSD. However, their experimental N treatments differ from the total soil N measurements reported here, which do not directly indicate plant-available N or vine N uptake. Similarly, other studies investigating plant nutrition have shown that NPK fertilization and high nutrient availability enhances vigour and therefore enhances foliar symptom expression (Calzarano *&* Di Marco, 2007; Calzarano *et al*., 2009). Gastou *et al*. (2024) reported that, at the cultivar level, lower vigor was associated with reduced GLSD foliar symptoms incidence. Consistently, vine vigor has been found to be positively correlated with the incidence of grapevine trunk diseases (GTDs) (Claverie *et al*., 2025).

Within this context, the greater root explorability observed in HRSVAs likely increases nutrient uptake (Koevoets *et al*., 2016), contributing to enhanced grapevine vigour (Gambetta *et al*., 2012). Among the nutrients, the one that differed markedly between LRSVAs and HRSVAs soil profiles is available manganese (Mn). Specifically, higher available Mn concentrations were measured in HRSVAs, particularly in the deeper horizons. This association raises the hypothesis that Mn may contribute to GLSD development. Recent *in vitro* studies by Bigazzi *et al*. (2026) demonstrated that Mn enhances wood degradation activity by *F. mediterranea*, one of the key fungi associated with ECDs. This pathogen is more frequently detected in symptomatic vines (Maher *et al*., 2012; Bruez *et al*., 2020), with increasing evidence supporting its central role in disease development and progression (Bruez *et al*., 2021b; Pacetti *et al*., 2021). In particular, *in vitro* experiments showed that Mn can increase manganese peroxidase activity, a major ligninolytic enzyme involved in wood decay (Bigazzi *et al*., 2026).

Indirect support for the involvement of Mn in wood degradation comes from Blanchette (1984), who investigated wood decayed by other white-rot fungi and reported markedly higher Mn concentrations in “black lines” or “black zones” surrounding decayed wood than in adjacent undecayed tissues, which showed little to no Mn accumulation. Similar dark zones have also been described in association with *F. mediterranea* (Mugnai *et al*., 1999).

In addition, soils in LRSVAs were characteriseds by a more alkaline pH compared to HRSVAs soils. Soil pH strongly influences Mn availability, which generally increases as pH decreases (Marschner, 2011). Furthermore, soil organic matter (SOM), which was higher in HRSVAs, can increase micronutrient availability by enhancing cation exchange capacity and forming soluble complexes that facilitate plant uptake (Weil & Brady, 2017; Dhaliwal *et al*., 2019). While these soil properties may help explain the differences in available Mn, they do not directly demonstrate greater Mn uptake or accumulation in the vines. We hypothesize that, if higher soil Mn availability translates into greater Mn accumulation within woody tissues, it could favour Mn-dependent ligninolytic activity by *F. mediterranea*, potentially contributing to wood degradation and, indirectly, GLSD symptom expression. However, since Mn concentrations in vine tissues and the extent of wood decay were not measured at these sites this proposed link remains hypothetical and requires targeted experimental testing.

Concluding, the comparison of soil properties between LRSVAs and HRSVAs suggests that GLSD symptoms expression is associated with contrasting edaphic conditions. HRSVAs were generally characterised by higher fertility, greater nutrient availability, and better root exploration ability, whereas LRSVAs showed lower fertility and physical limitations related to soil drainage, including temporary waterlogging conditions. Importantly, despite these differences, the two soil profiles in areas with the same disease recurrence class (high or low), displayed consistent soil characteristics, regardless of whether they had been excavated in areas characterized by different levels of ECa. This suggest that distinct recurring soil conditions are associated with a varying probability of GLSD expression. Nevertheless, further studies across multiple vineyards are needed to validate this hypothesis. Notably, the identification of such clear patterns within a relatively small study area further highlights the inherent spatial heterogeneity of vineyard soils and underscores the need for more comprehensive investigations to better disentangle the role of abiotic factors in GLSD expression, beyond climatic drivers such as rainfall.

## Conclusions

In this case study, we showed that GLSD symptom expression is associated with the spatial heterogeneity of soil physical, hydrological and chemical features, which is reflected in a structured spatial pattern of disease distribution. These findings suggest that soil act as an important abiotic driver of foliar symptoms development in GLSD, the most common disease within the esca complex of diseases, with both macro- and micronutrients potentially contributing to disease expression. Contrary to what one might expect, limitations to rooting depth, such as temporary waterlogging and pretty high active lime content, provide light abiotic stress and plant vigour limitation, but concurrently reduces the recurrence of GLSD symptoms. Conversely, deep and fertile soils were associated with a higher recurrence of GLSD symptoms. In addition, higher content of available Mn in areas with higher GLSD recurrence, merits to be further investigated, also taking into account its ability to enhance wood degradation activity by *F. mediterranea*, one of the major fungi associated with esca complex of diseases, the one that actually gave the disease its name (Graniti, 2006).

The identification of structured spatial patterns of disease distribution, together with the marked edaphic heterogeneity observed within a single vineyard, highlights the importance of investigating soil–plant interactions at a finer spatial scale. Although this preliminary results require validation across additional sites, they provide further evidence that soil deserves greater attention as an abiotic factor influencing GLSD, a factor that has remained underexplored despite growing evidence of its role in disease development and symptoms expression.

## Acknowledgements

Special thanks to Prof Giuseppe Surico for his critical reading of the manuscript and constructive suggestions.

## Notes

### Competing Interest Statement

The authors have declared no competing interest.

